## Supplementary Information for "Fluorescent protein lifetimes report increased local densities and phases of nuclear condensates during embryonic stem cell differentiation"

[*] Shared first authors

[‡] Corresponding authors

**Supplementary Information and Equations**

Retrieving the fluorescence lifetime values from fitting models to fluorescence decays

Theoretical models fitted to fluorescence decays (Eq. 1):

$$I(t) \propto irf(t) \otimes \left(\alpha_1 e^{-\frac{t}{\tau_1}} + \alpha_2 e^{-\frac{t}{\tau_2}}\right) \quad (1)$$

where $t$ is photon nanotime, the time between the moment of excitation and the moment of photon detection, $I(t)$ is the photon counts as a function of photon nanotime, $irf(t)$ is the impulse response function (IRF) of the system, and $\tau_1$, $\tau_2$, and $\alpha_1$, $\alpha_2$, are the two lifetime components and their amplitudes, respectively. The $\otimes$ operator is a commonly used symbol of the convolution operation. Then, the two lifetime components are used for the calculation of the intrinsic average lifetime, $\bar{\tau}$ (Eq. 2):

$$\bar{\tau} = \frac{\alpha_1\tau_1^2 + \alpha_2\tau_2^2}{\alpha_1\tau_1 + \alpha_2\tau_2} \quad (2)$$

Retrieving the fluorescence anisotropy decay curves from polarized fluorescence decays

The fluorescence anisotropy decay, $r(t)$, was calculated from the polarized fluorescence decays as follows (Eq. 3):

$$r(t) = \frac{I_{\parallel}(t) - G \cdot I_{\perp}(t)}{I_{\parallel}(t) + 2G \cdot I_{\perp}(t)} \quad (3)$$

where $I_{\parallel}(t)$ and $I_{\perp}(t)$ are the decays of fluorescence polarized parallel- and perpendicular relative to the excitation polarization, $t$ is the time following the moment of excitation, and $G$ is the well-known G-factor which represents deviations from equality of $I_{\parallel}$ and $I_{\perp}$ in samples not exhibiting fluorescence anisotropy.

The fluorescence anisotropy due to a given depolarization mode, $r$, depends both on the fluorescence lifetime, $\tau$, and the rotational correlation time, $\theta$, following the Perrin equation (Eq. 4):

$$\frac{r_0}{r} = 1 + \frac{\tau}{\theta} \quad (4)$$

Where, $r_0$, the fundamental anisotropy, is a constant for a given dye system. Importantly, the rotational correlation time is influenced either by temperature, viscosity or size of the rotating system, following the Stokes-Einstein relationship, or due to faster depolarization that can be caused by energy transfer.

In-cell FCS calculations

The fluorescence autocorrelation curves, $G(\tau)$, were calculated from the acquired fluorescence intensity trajectories, $I(t)$ (Eq. 5):

$$G(\tau) = \frac{\langle \delta I(t) \cdot \delta I(t+\tau) \rangle}{\langle I(t) \rangle^2} \quad (5)$$

where $t$ is the absolute photon arrival time, $\tau$ is the time lag between a pair of photon arrival times, $\delta$ denotes the deviation relative to the mean value and $\langle\ \rangle$ denote a time average.

Then, the resulting fluorescence autocorrelation curves were fitted to a model that takes into account fluorescence fluctuations due to the diffusion of two species in and out of the laser focus, fast and slower diffusing species (Eq. 6):

$$G(\tau) = \frac{1}{\bar{N}} \cdot \left\{ \frac{f}{\left(1+\frac{\tau}{\tau_{D,fast}}\right)\sqrt{1+\left(\frac{\omega_0}{z_0}\right)^2 \cdot \frac{\tau}{\tau_{D,fast}}}} + \frac{1-f}{\left(1+\frac{\tau}{\tau_{D,slow}}\right)\sqrt{1+\left(\frac{\omega_0}{z_0}\right)^2 \cdot \frac{\tau}{\tau_{D,slow}}}} \right\} \quad (6)$$

where $\bar{N}$ is the mean number of molecules in the laser focus, f is the fraction of the fast species, $\tau_{D,fast}$ and $\tau_{D,slow}$ are the diffusion times through the laser focus of the fast and slow diffusing species, respectively, and $\omega_0$ and $z_0$ are the axial and longitudal lengths of the laser focus. Fitting was performed after constraining the model parameters so that all parameters should have positive values, the fraction parameter should not exceed a value of 1 and the ratio of the longitudinal and axial lengths of the laser focus should be bound within the typical 4 – 10 range of values.

Analyses of photobleaching rates

The acquired in-cell fluorescence trajectories are exhibiting two timescales of photobleaching, slow and fast, representing rapidly and slowly diffusing species. Therefore, each in-cell fluorescence intensity trajectory was fitted with a bi-exponential decay model (Eq. 7):

$$I(t) = a_{fast}e^{-\frac{t}{t_{fast}}} + a_{slow}e^{-\frac{t}{t_{slow}}} + y_0 \quad (7)$$

where $t_{fast}$ and $t_{slow}$ are the lifetimes of the fast and slow photobleaching process, $a_{fast}$ and $a_{slow}$ are their amplitude and $y_0$ is a baseline representing species that diffuse fast enough that they do not photobleach even within the 2 minutes of acquisition.

### Supplementary Figures

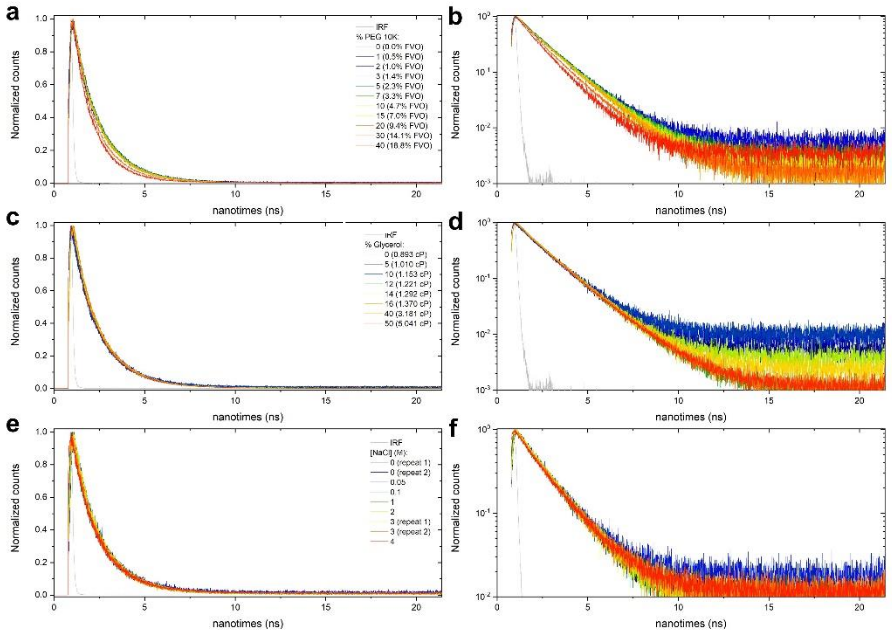

**Supplementary Fig. 1: mCherry normalized fluorescence decays at different conditions. (a, c, e)** linear scale. **(b, d, f**) log-scale. **(a, b)**, fluorescence decays of 100 nM mCherry as a function of PEG 10,000; **(c, d)** decays as a function of glycerol; **(e, f)** decays as a function of NaCl.

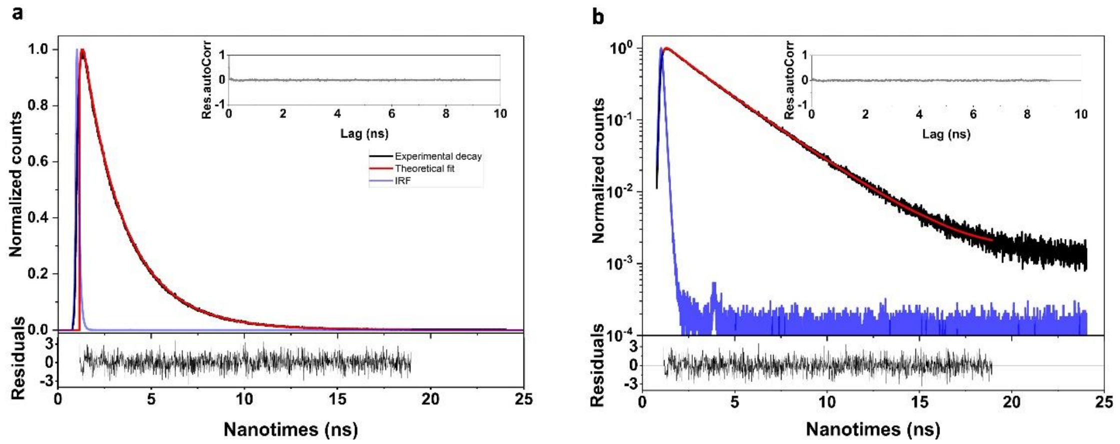

**Supplementary Fig. 2: An example of the fluorescence decay fitting procedure.** The fluorescence decay fitting is shown in linear **a**, and log scales **b**, respectively. Experimental decay (black), the IRF (blue) and the best fit model (red) are shown. The fitting residuals (bottom panels) and their auto-correlation (panel insets) are also shown. This example is the best-fit of the fluorescence decay of eGFP in the presence of 1.3 M trehalose, with reduced $\chi^2$ of 1.029.

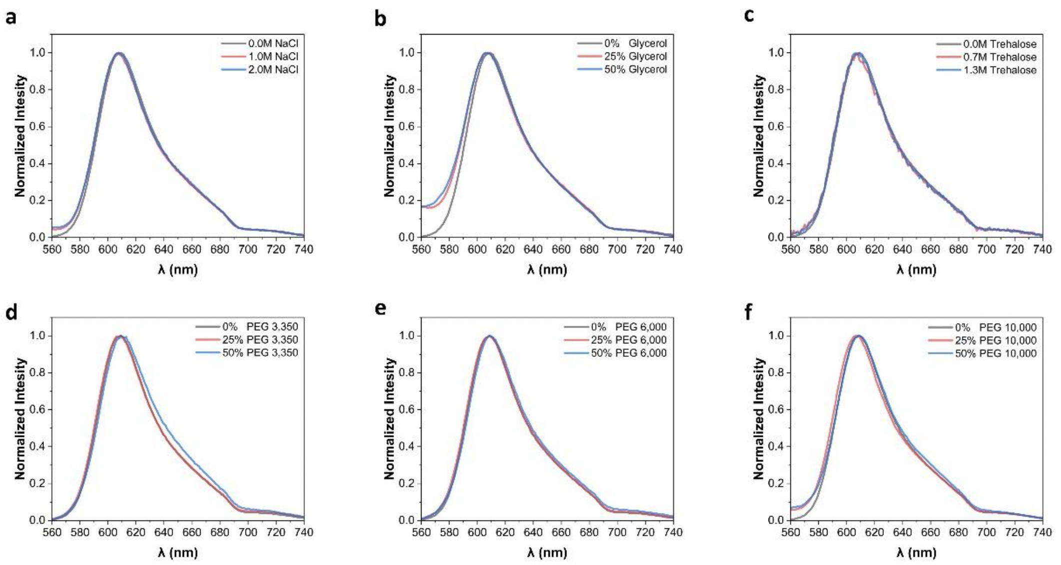

**Supplementary Fig. 3: mCherry normalized fluorescence spectra at different conditions.** The fluorescence spectra of 100 nM mCherry were measured at different concentrations of different conditions: **a**, NaCl, **b**, Glycerol, **c**, Trehalose, **d**, PEG 3,350, **e**, PEG 6,000 and **f**, PEG 10,000.

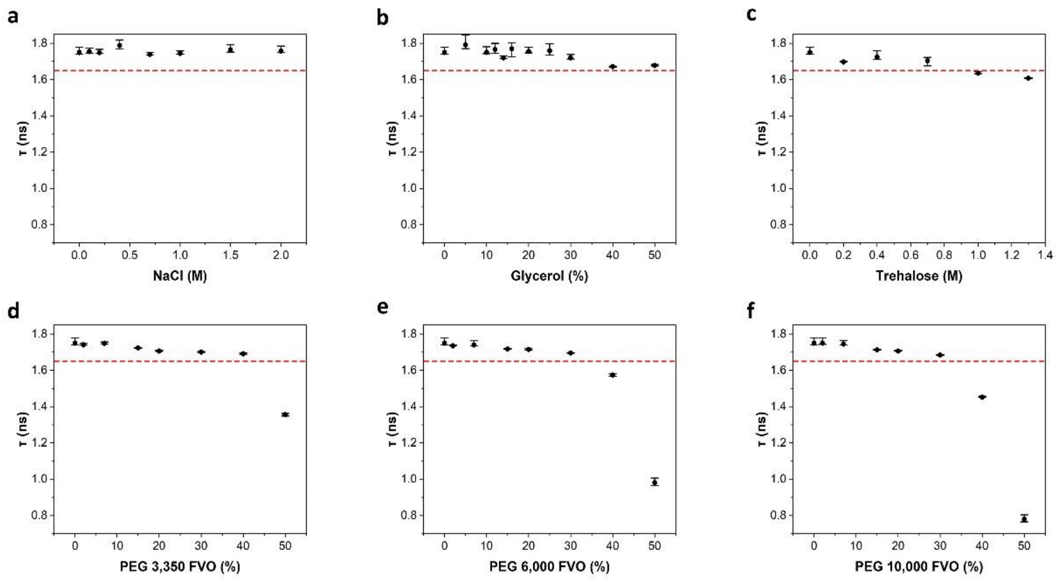

**Supplementary Fig. 4: The mean fluorescence lifetime of mRFP at different physico-chemical conditions.** mean fluorescence lifetime (τ) of 100 nM mRFP as a measure of increasing NaCl concentrations **a**, Glycerol, **b**, Trehalose, **c**, and **d-f**, PEG. Crowding exclusively induces lifetime reduction, and only above 35-45% FVO. The values and error estimates are based on calculations (Eq. S2) using best-fit values of the recorded fluorescence decays (Fig. S1) to a sum of two exponents function (Eq. S1). The values of the intrinsic mean fluorescence lifetimes are reported in Table S1. The error estimates are the minimal and maximal intrinsic mean fluorescence lifetime values calculated from all lifetime component values and their amplitudes, which are within a reduced $\chi^2$ that is within 95% confidence relative to the minimal best-fit reduced $\chi^2$ value. Red line that is set at 1.65 ns highlights the limit below which fluorescence lifetimes drop in the presence of elevated concentrations of PEG or trehalose. Titration of NaCl or glycerol, leads to non-monotonic changes in τ which fluctuate around the typical fluorescence lifetime of mRFP.

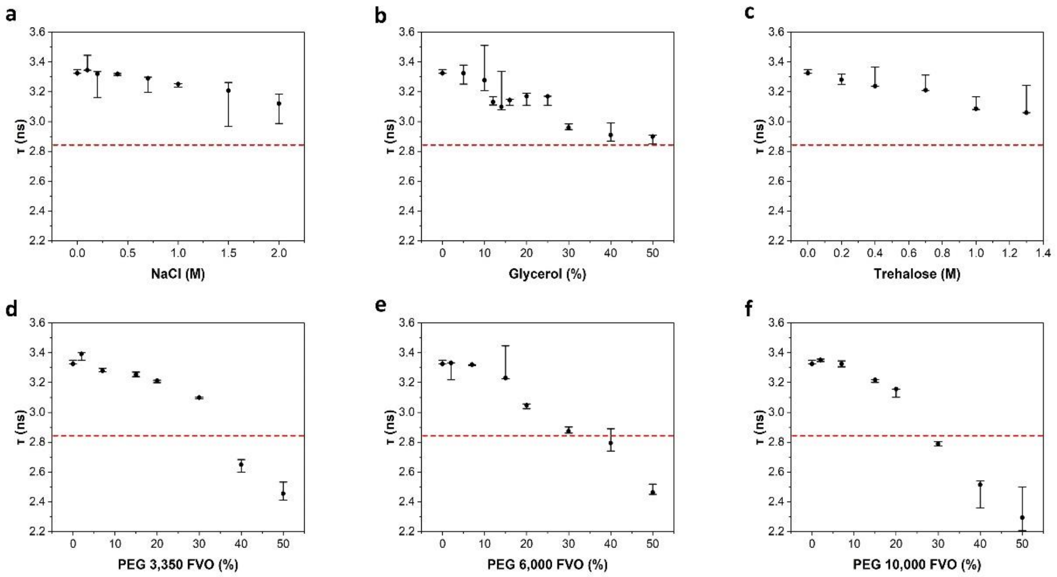

**Supplementary Fig. 5: The mean fluorescence lifetime of mCitrine at different physico-chemical conditions.** mean fluorescence lifetime (τ) of 100 nM mCitrine as a measure of increasing NaCl concentrations **a**, Glycerol, **b**, Trehalose, **c**, and **d-f**, PEG. Crowding exclusively induces lifetime reduction, and only above 30% FVO. The values and error estimates are based on calculations (Eq. S2) using best-fit values of the recorded fluorescence decays (Fig. S1) to a sum of two exponents function (Eq. S1). The values of the intrinsic mean fluorescence lifetimes are reported in Table S1. The error estimates are the minimal and maximal intrinsic mean fluorescence lifetime values calculated from all lifetime component values and their amplitudes, which are within a reduced $\chi^2$ that is within 95% confidence relative to the minimal best-fit reduced $\chi^2$ value. Red line that is set at 2.84 ns highlights the limit below which fluorescence lifetimes drop in the presence of elevated concentrations of PEG or trehalose.

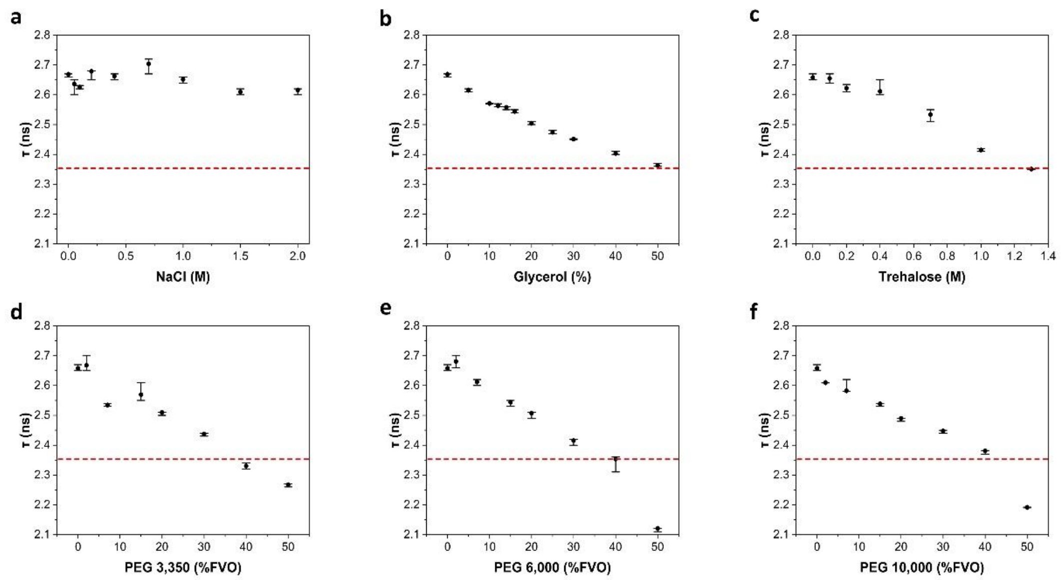

**Supplementary Fig. 6: The mean fluorescence lifetime values of eGFP at different concentrations of different physico-chemical conditions.** mean fluorescence lifetime (τ) of 100 nM eGFP as a measure of increasing NaCl concentrations **a**, Glycerol, **b**, Trehalose, **c**, and **d-f**, PEG. Viscosity and crowding induce lifetime reduction. Fluorescence lifetime reduction from the typical 2.5-2.7 ns is induced above 40% FVO. The values of the intrinsic mean fluorescence lifetimes are reported in Table S1. The error estimates are the minimal and maximal intrinsic mean fluorescence lifetime values calculated from all lifetime component values and their amplitudes, which are within a reduced $\chi^2$ that is within 95% confidence relative to the minimal best-fit reduced $\chi^2$ value. Red line that is set at 2.35 ns highlights the limit below which fluorescence lifetimes drop in the presence of elevated concentrations of PEG or trehalose.

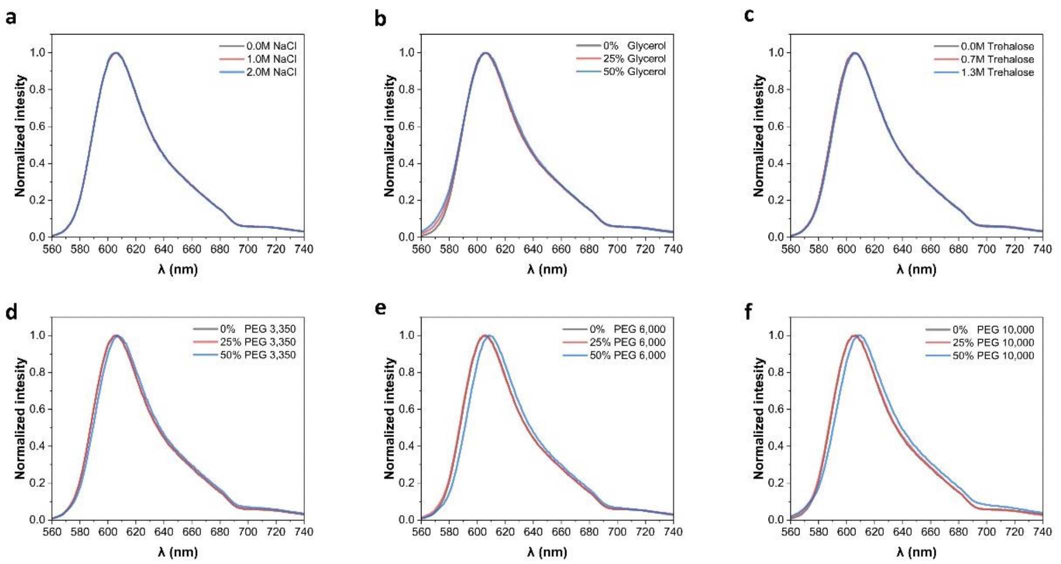

**Supplementary Fig. 7: mRFP normalized fluorescence spectra at different conditions.** The fluorescence spectra of 100 nM mRFP were measured at different concentrations of different conditions: **a**, NaCl, **b**, Glycerol, **c**, Trehalose, **d**, PEG 3,350, **e**, PEG 6,000 and **f**, PEG 10,000.

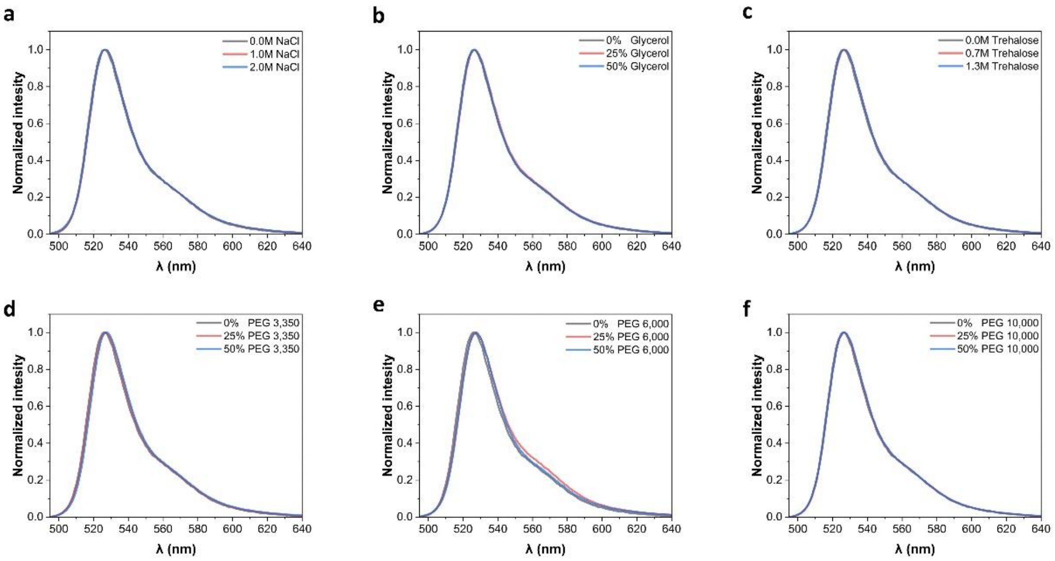

**Supplementary Fig. 8: mCitrine normalized fluorescence spectra at different conditions.** The fluorescence spectra of 100 nM mCitrine were measured at different concentrations of different conditions: **a**, NaCl, **b**, Glycerol, **c**, Trehalose, **d**, PEG 3,350, **e**, PEG 6,000 and **f**, PEG 10,000.

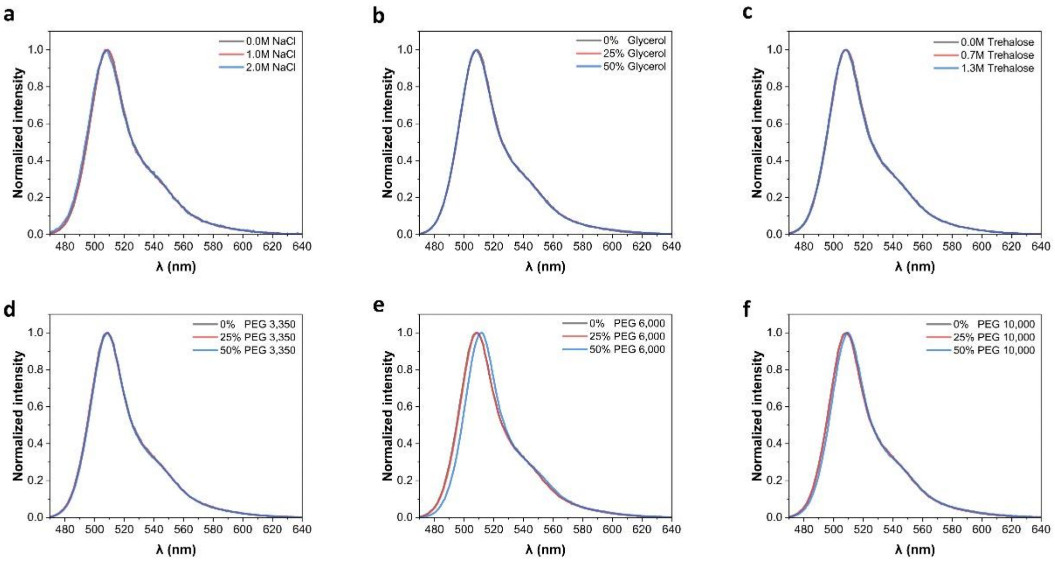

**Supplementary Fig. 9: eGFP normalized fluorescence spectra at different conditions.** The fluorescence spectra of 100 nM eGFP were measured at different concentrations of different conditions: **a**, NaCl, **b**, Glycerol, **c**, Trehalose, **d**, PEG 3,350, **e**, PEG 6,000 and **f**, PEG 10,000.

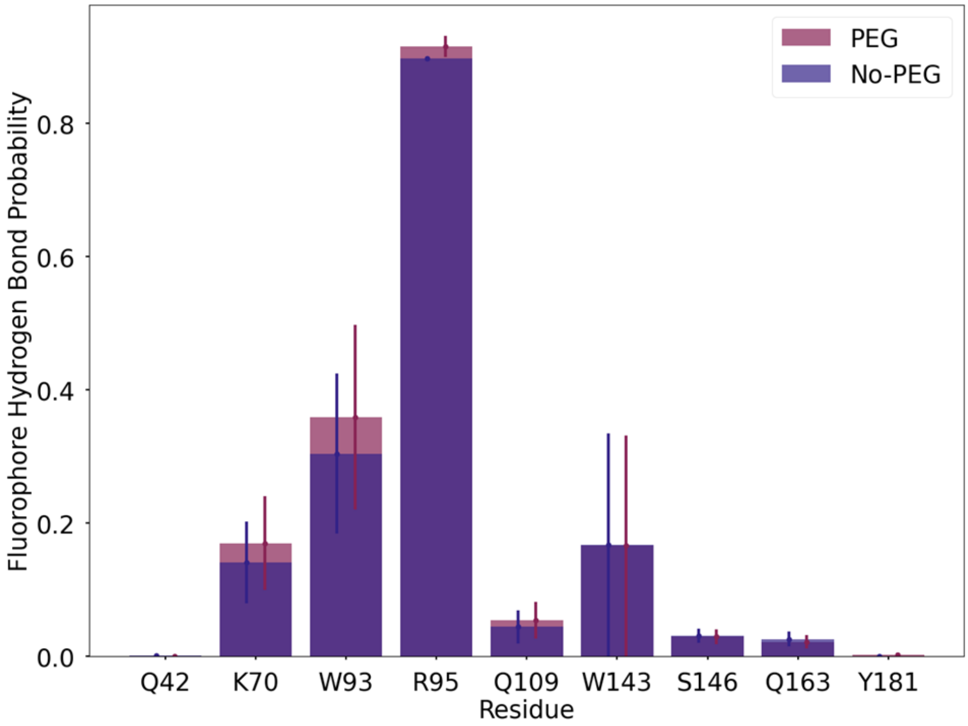

**Supplementary Fig. 10: Protein-fluorophore hydrogen bonds.** The probability of hydrogen bond formation between the fluorophore and other protein residues for each system. Values for the PEG system are shown in plum and values for the no-PEG system are shown in navy blue. Uncertainty is shown as error bars, computed as the standard error of the mean treating each trajectory as an independent measurement.

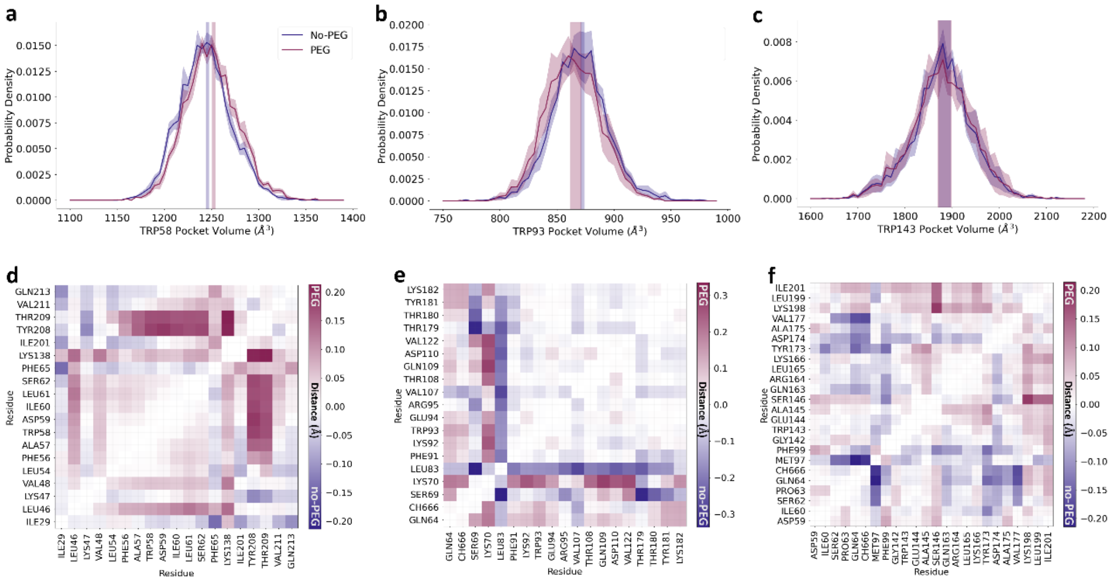

**Supplementary Fig. 11: Trp environments**. **a-c** show the distributions of Trp pocket volumes for Trp's 58, 93, and 143, respectively. The distributions for the PEG systems are shown in plum, and the distributions for the systems without PEG are shown in navy. Shaded regions indicate the standard error of the mean, treating each trajectory as an independent observation. **d-f** show the difference map of the average Trp-pocket $C_\alpha$-$C_\alpha$ distances for Trp's 58, 93, and 143, respectively. Distances that are larger in the PEG system are positive and shown in plum, and ones that are larger in the no-PEG system are negative and shown in navy.

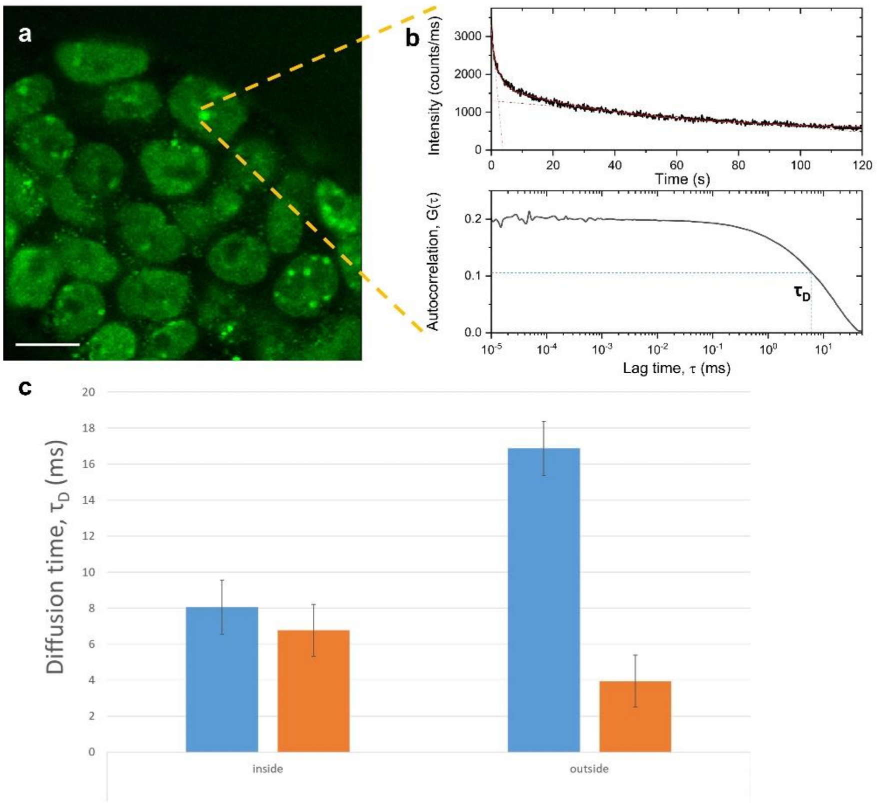

**Supplementary Fig. 12: In-cell FCS report on diffusivities within HP1α condensates. a,** HP1α-mCherry image shows nuclei with bright foci. We position the laser focus onto a given position within one such HP1α-mCherry condensate (yellow dashed arrows), and acquire fluorescence for a given acquisition time (e.g., 2 minutes). **b,** The fluorescence trajectory acquired (top) from the chosen position within the HP1α-mCherry condensate exhibits a monotonic decrease due to photobleaching. This intensity trajectory is fitted to a bi-exponential decay to characterize the photobleaching rates. The fluorescence trajectory is noisy, and this noise carries information regarding the temporal fluctuations in the number of molecules within the laser focus and the diffusion times of the mCherry-tagged HP1α molecules traversing the laser focus. The fluctuations and diffusion times are retrieved from the autocorrelation (bottom panel) of the fluorescence trajectory. The characteristic diffusion time, $\tau_D$ (blue dashed line), is retrieved by model fitting (Eq. 4) to the fluorescence autocorrelation curve (Eq. S3). Of note is that the timescales covered by the fluorescence autocorrelation curve (bottom; <100 ms) are faster than the two photobleaching decay times observed in the acquired fluorescence trajectory (top; seconds and tens of seconds). **c,** Averages of diffusion times inside (left) and outside (right) multiple HP1α-mCherry condensates before (blue) and after (orange) early differentiation are reported. Despite the reported differences in diffusion times, the general conclusion is that diffusion of HP1α-mCherry occurs within condensates under all conditions tested, and hence a solid phase can be ruled-out.

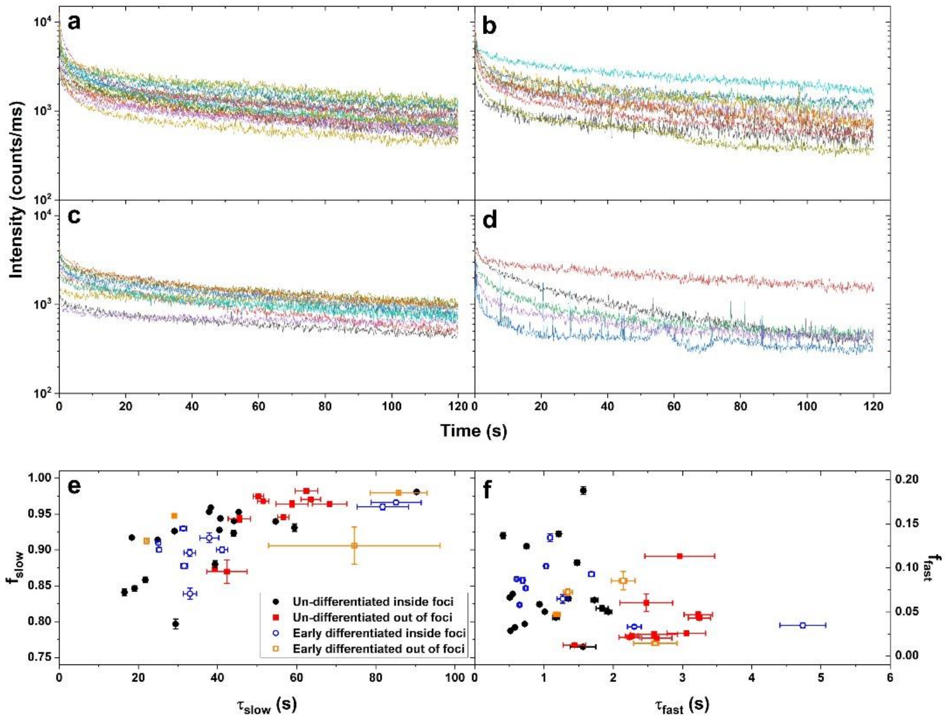

**Supplementary Fig. 13: Photobleaching rates analyses of in-cell fluorescence intensity trajectories. a-d**, Multiple 2-minute fluorescence intensity trajectories were recorded inside (**a**, **b**) and outside (**c**, **d**) of HP1α-mCherry foci, in un-differentiated (**a**, **c**) and early differentiated (**b**, **d**) ESCs. **e, f**, summarizes the fractions of the slow and fast photobleaching processes, and their times inside HP1α-mCherry foci and out-of-foci in un-differentiated and in early differentiated ESCs.

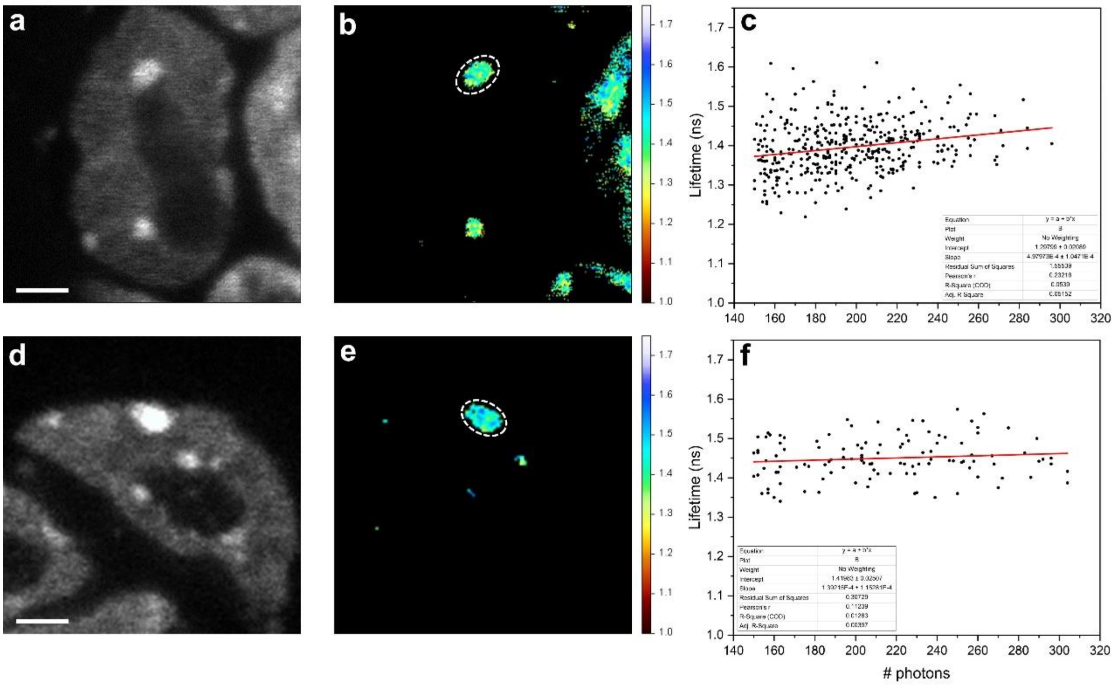

**Supplementary Fig. 14: Fluorescence intensities and lifetimes within HP1α condensates exhibit at most weak correlation. a-f,** Two examples of undifferentiated ESCs showing HP1α condensates (**a**, **b** and **d**, **e**) with heterogeneity in fluorescence lifetimes (**b**, **e**). The Pearson correlation coefficient values between pixel intensity (**a**, **d**) and lifetime (**b**, **e**) are 0.23 (**c**) and 0.11 (**f**), pointing towards a weak correlation at most between changes in intensity and changes in lifetimes (**c**, **f**). Scale bar 2 µm. Fluorescence lifetime color bars scale from 1.00 to 1.75 ns.

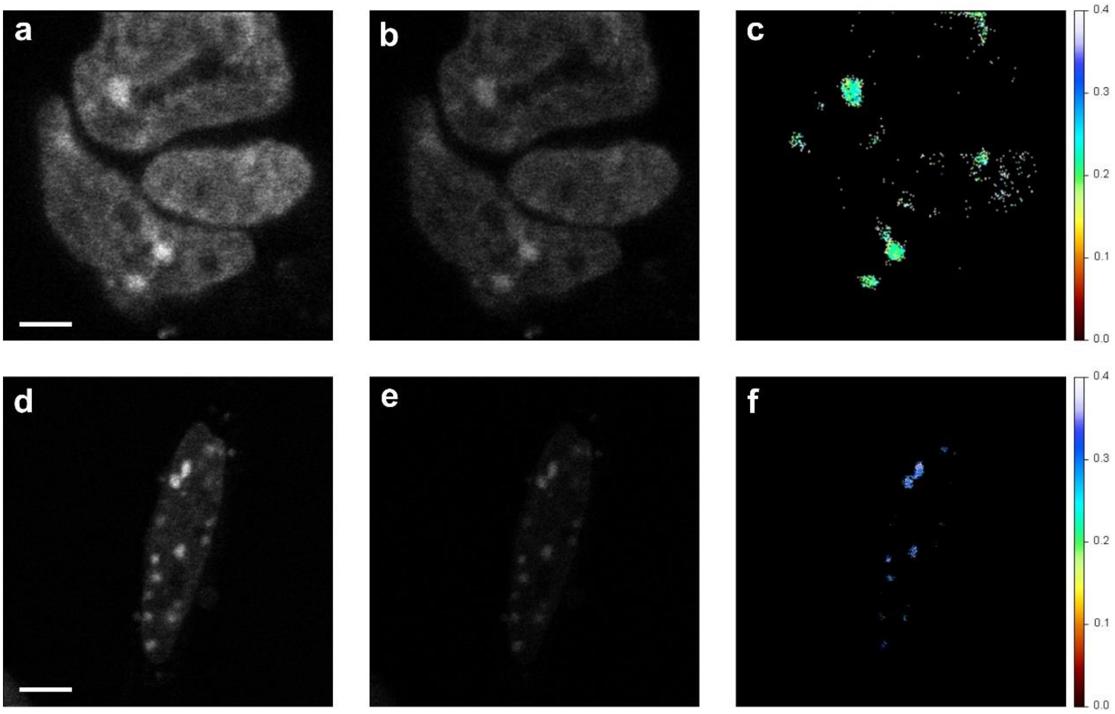

**Supplementary Fig. 15: Fluorescence anisotropy images recover differences between HP1α condensates in undifferentiated and early-differentiated ESCs. a**, **b**, **d**, **e**, fluorescence intensity images of fluorescence polarized parallel (**a**, **d**) and perpendicular (**b**, **e**) relative to the polarization of excitation, in undifferentiated (**a**, **b**) and early-differentiated (**d**, **e**) cells. **c**, **f,** the calculated fluorescence anisotropy images for pixels within HP1α condensates, in undifferentiated (**c**) and early-differentiated (**f**) cells. Scale bar 2 µm. Fluorescence anisotropy color bars scale from 0.0 to 0.4 ns.

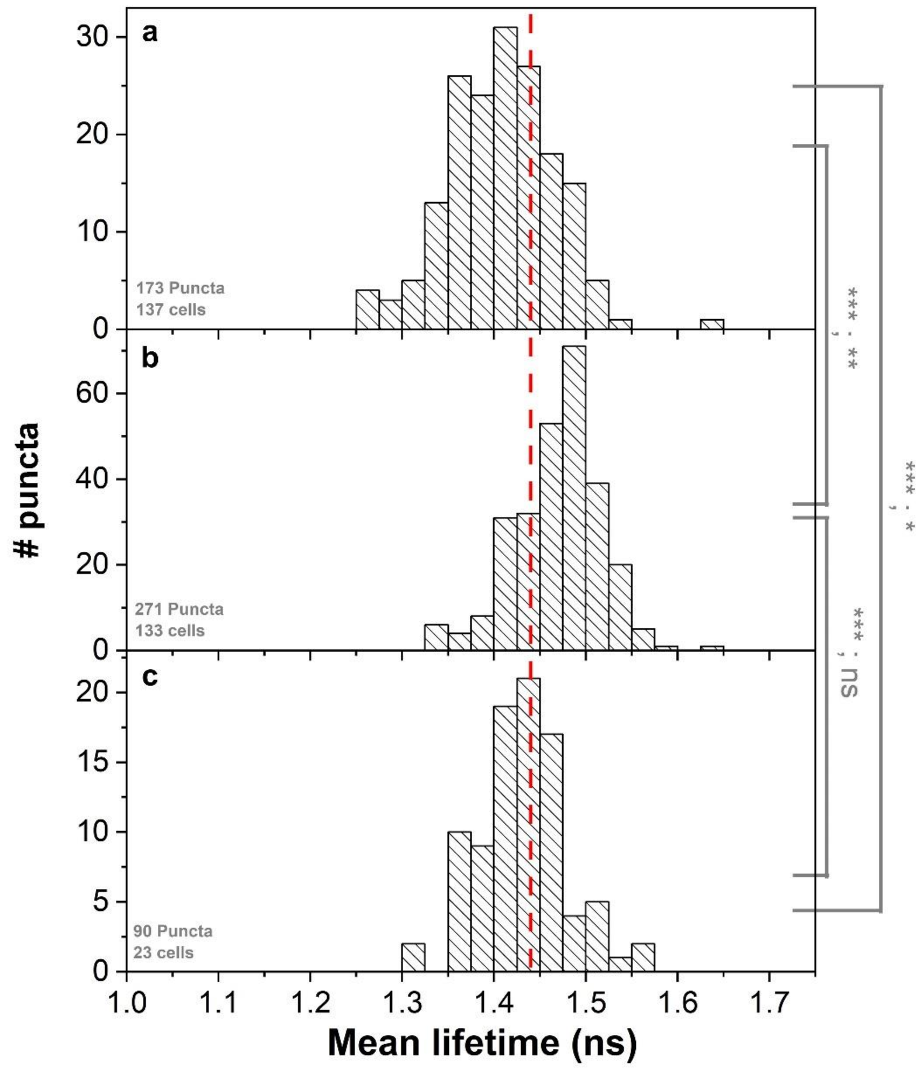

**Supplementary Fig. 16: Mean fluorescence lifetimes of mCherry-HP1α condensates from undifferentiated ESCs, early-differentiated ESCs and ESCs transfected with LNA probes.** **a,** Undifferentiated ESCs; **b,** Early-differentiated ESCs and **c,** Undifferentiated ESCs transfected with LNA probes. Histograms show the distribution of mean fluorescence lifetimes of different mCherry-HP1α puncta. Asterisks report on the p-value from two-sided T-test and from two-sided F-test, comparing means and variances of histograms. $p>0.05$ n.s.; $0.01<p<0.05^{*}$; $0.001<p<0.01^{**}$; $p<0.001^{***}$. A vertical red line shows the 1.44 ns lower boundary of mCherry fluorescence lifetime below which the crowdedness level is above 30% FVO (Fig. 1).

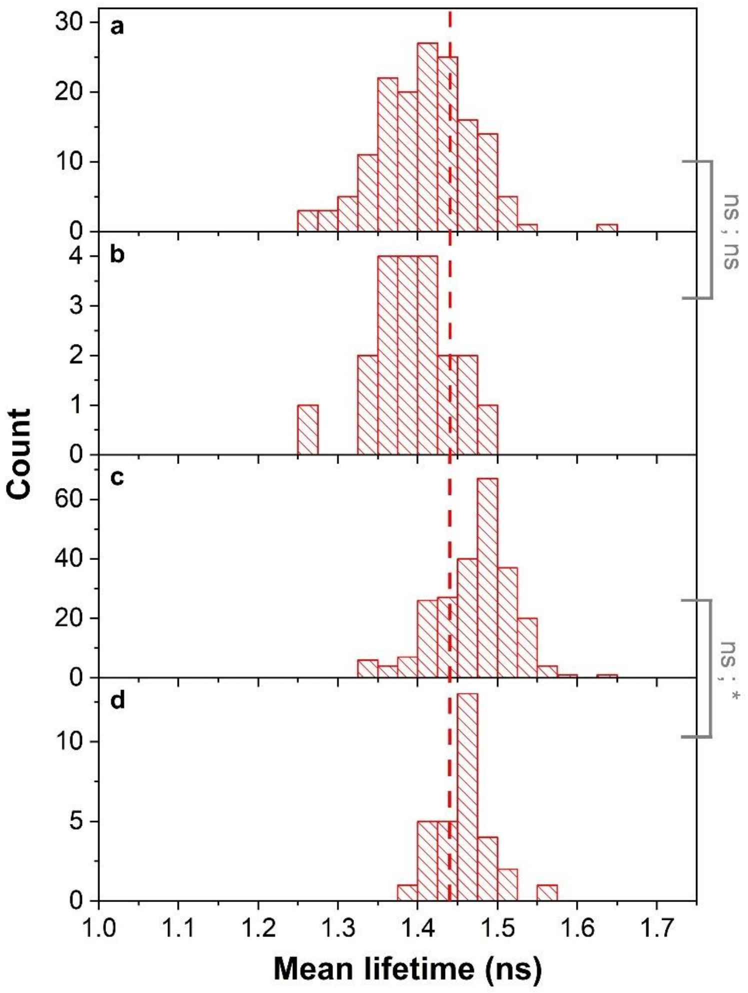

**Supplementary Fig. 17: Mean fluorescence lifetimes in mCherry-HP1α condensates from undifferentiated and early-differentiated ESCs in the absence and presence of transfected RNA scramble probe.** **a**, **b**, A comparison of the mean fluorescence lifetime histograms in undifferentiated ESCs in the absence (**a**) and presence (**b**) of transfected RNA scramble probe, showing lack of significant differences, allowing their summation. **c**, **d**, A comparison of the mean fluorescence lifetime histograms in early-differentiated ESCs in the absence (**c**) and presence (**d**) of transfected RNA scramble probe, showing lack of significant differences, allowing their summation. Asterisks report on the p-value from two-sided T-test and from two-sided F-test, comparing means and variances of histograms. $p>0.05$ n.s.; $0.01<p<0.05^{*}$; $0.001<p<0.01^{**}$; $p<0.001^{***}$. A vertical red line shows the 1.44 ns lower boundary of mCherry fluorescence lifetime below which the crowdedness level is above 30% FVO (Fig. 1).

### Supplementary Tables

**Supplementary Table 1**: best fit results of fluorescence decay fittings and intrinsic average lifetimes. The error estimates are the minimal and maximal intrinsic mean fluorescence lifetime values calculated from all lifetime component values and their amplitudes, which are within a reduced $\chi^2$ that is within 95% confidence relative to the minimal best-fit reduced $\chi^2$ value.

| FP | Additive | Concentration (M)/Viscosity (% glycerol)/Crowding (%FVO) | $\alpha_1$ | $\tau_1$ (ns) | $\alpha_2$ | $\tau_2$ (ns) | $\bar{\tau}$ (ns) | $\chi_R^2$ |
|---|---|---|---|---|---|---|---|---|
| mCherry | NaCl | 0.0 (M) | 4.33 | 1.59 | 1.59 | 0.83 | 1.46 (1.46-1.47) | 1.240 |
| mCherry | NaCl | 0.05 (M) | 5.70 | 1.55 | 1.48 | 0.74 | 1.47 (1.46-1.47) | 1.216 |
| mCherry | NaCl | 0.1 (M) | 5.94 | 1.45 | 0.35 | 1.45 | 1.45 (1.44-1.46) | 1.085 |
| mCherry | NaCl | 0.2 (M) | 5.94 | 1.50 | 1.04 | 0.48 | 1.45 (1.44-1.45) | 1.077 |
| mCherry | NaCl | 0.4 (M) | 5.72 | 1.50 | 1.00 | 0.50 | 1.44 (1.44-1.45) | 1.135 |
| mCherry | NaCl | 0.7 (M) | 5.90 | 1.50 | 1.00 | 0.52 | 1.45 (1.44-1.46) | 1.045 |
| mCherry | NaCl | 1.0 (M) | 1.97 | 1.18 | 0.95 | 1.91 | 1.50 (1.49-1.52) | 1.092 |
| mCherry | NaCl | 1.5 (M) | 0.91 | 0.82 | 2.43 | 1.59 | 1.47 (1.46-1.48) | 1.113 |
| mCherry | NaCl | 2.0 (M) | 1.16 | 0.95 | 1.99 | 1.66 | 1.48 (1.47-1.49) | 1.174 |
| mCherry | Trehalose | 0.0 (M) | 2.35 | 1.17 | 1.41 | 1.87 | 1.52 (1.50-1.53) | 1.275 |
| mCherry | Trehalose | 0.2 (M) | 0.71 | 0.78 | 2.47 | 1.59 | 1.49 (1.48-1.50) | 1.179 |
| mCherry | Trehalose | 0.4 (M) | 0.81 | 0.78 | 2.89 | 1.58 | 1.49 (1.48-1.49) | 1.196 |
| mCherry | Trehalose | 0.7 (M) | 3.93 | 1.53 | 0.73 | 0.56 | 1.46 (1.45-1.47) | 1.165 |
| mCherry | Trehalose | 1.0 (M) | 0.83 | 0.43 | 4.69 | 1.48 | 1.43 (1.42-1.43) | 1.110 |
| mCherry | Trehalose | 1.3 (M) | 6.58 | 1.39 | 1.47 | 0.34 | 1.34 (1.33-1.35) | 1.091 |
| mCherry | Glycerol | 0 (%) | 6.31 | 1.47 | - | - | 1.47 (1.45-1.48) | 1.158 |
| mCherry | Glycerol | 5 (%) | 3.50 | 1.58 | 0.82 | 0.76 | 1.50 (1.48-1.51) | 1.152 |
| mCherry | Glycerol | 10 (%) | 4.23 | 1.53 | 0.67 | 0.54 | 1.48 (1.47-1.49) | 1.075 |
| mCherry | Glycerol | 12 (%) | 1.01 | 0.98 | 2.24 | 1.65 | 1.51 (1.49-1.52) | 1.129 |
| mCherry | Glycerol | 14 (%) | 7.31 | 1.54 | 1.14 | 0.48 | 1.49 (1.48-1.49) | 1.093 |
| mCherry | Glycerol | 16 (%) | 4.11 | 1.56 | 0.84 | 0.66 | 1.49 (1.48-1.50) | 1.127 |
| mCherry | Glycerol | 20 (%) | 1.05 | 0.35 | 7.53 | 1.51 | 1.47 (1.46-1.48) | 1.103 |
| mCherry | Glycerol | 25 (%) | 4.47 | 1.31 | 3.54 | 1.67 | 1.49 (1.47-1.49) | 1.116 |
| mCherry | Glycerol | 30 (%) | 8.34 | 1.52 | 0.87 | 0.47 | 1.48 (1.48-1.49) | 1.101 |
| mCherry | Glycerol | 40 (%) | 3.66 | 1.68 | 6.05 | 1.36 | 1.49 (1.48-1.49) | 1.108 |
| mCherry | Glycerol | 50 (%) | 5.83 | 1.57 | 1.22 | 0.54 | 1.50 (1.49-1.51) | 1.102 |
| mCherry | PEG 3,350 | 0 (% FVO) | 0.99 | 0.66 | 6.62 | 1.55 | 1.50 (1.49-1.50) | 1.586 |
| mCherry | PEG 3,350 | 2 (% FVO) | 5.08 | 1.52 | 0.66 | 0.54 | 1.48 (1.47-1.49) | 1.204 |
| mCherry | PEG 3,350 | 7 (% FVO) | 5.90 | 1.57 | 1.50 | 0.76 | 1.48 (1.48-1.49) | 1.161 |
| mCherry | PEG 3,350 | 15 (% FVO) | 9.58 | 1.50 | 1.45 | 0.36 | 1.46 (1.45-1.47) | 1.168 |
| mCherry | PEG 3,350 | 20 (% FVO) | 12.40 | 1.49 | 1.87 | 0.33 | 1.45 (1.44-1.46) | 1.219 |
| mCherry | PEG 3,350 | 30 (% FVO) | 11.72 | 1.50 | 2.27 | 0.52 | 1.44 (1.43-1.45) | 1.125 |
| mCherry | PEG 3,350 | 40 (% FVO) | 8.87 | 1.38 | 2.07 | 0.44 | 1.32 (1.31-1.32) | 1.178 |
| mCherry | PEG 3,350 | 50 (% FVO) | 6.55 | 1.39 | 2.09 | 0.54 | 1.30 (1.29-1.30) | 1.184 |
| mCherry | PEG 6,000 | 0 (% FVO) | 4.34 | 1.59 | 1.59 | 0.83 | 1.46 (1.46-1.47) | 1.240 |
| mCherry | PEG 6,000 | 2 (% FVO) | 5.81 | 1.51 | 0.96 | 0.52 | 1.46 (1.45-1.46) | 1.136 |
| mCherry | PEG 6,000 | 7 (% FVO) | 5.74 | 1.50 | 0.99 | 0.52 | 1.45 (1.44-1.45) | 1.137 |
| mCherry | PEG 6,000 | 15 (% FVO) | 8.31 | 1.49 | 1.78 | 0.46 | 1.44 (1.43-1.45) | 1.201 |
| mCherry | PEG 6,000 | 20 (% FVO) | 8.45 | 1.49 | 1.57 | 0.44 | 1.44 (1.43-1.45) | 1.223 |
| mCherry | PEG 6,000 | 30 (% FVO) | 8.90 | 1.50 | 1.82 | 0.52 | 1.44 (1.43-1.45) | 1.210 |
| mCherry | PEG 6,000 | 40 (% FVO) | 4.49 | 1.49 | 2.17 | 0.66 | 1.35 (1.34-1.35) | 1.158 |
| mCherry | PEG 6,000 | 50 (% FVO) | 4.86 | 1.27 | 1.95 | 0.54 | 1.17 (1.16-1.17) | 1.203 |
| mCherry | PEG 10,000 | 0 (% FVO) | 4.34 | 1.59 | 1.59 | 0.83 | 1.46 (1.46-1.47) | 1.298 |
| mCherry | PEG 10,000 | 2 (% FVO) | 6.01 | 1.51 | 0.98 | 0.49 | 1.46 (1.45-1.46) | 1.110 |
| mCherry | PEG 10,000 | 7 (% FVO) | 8.29 | 1.49 | 1.12 | 0.40 | 1.45 (1.44-1.46) | 1.147 |
| mCherry | PEG 10,000 | 15 (% FVO) | 7.59 | 1.50 | 1.28 | 0.46 | 1.45 (1.44-1.45) | 1.194 |
| mCherry | PEG 10,000 | 20 (% FVO) | 9.46 | 1.50 | 1.78 | 0.46 | 1.44 (1.43-1.45) | 1.161 |
| mCherry | PEG 10,000 | 30 (% FVO) | 6.97 | 1.50 | 2.05 | 0.55 | 1.41 (1.40-1.42) | 1.176 |
| mCherry | PEG 10,000 | 40 (% FVO) | 6.14 | 1.45 | 4.13 | 0.61 | 1.27 (1.26-1.27) | 1.153 |
| mCherry | PEG 10,000 | 50 (% FVO) | 2.66 | 0.92 | 1.53 | 1.61 | 1.26 (1.25-1.27) | 1.192 |
| mRFP | NaCl | 0.0 (M) | 3.21 | 1.50 | 0.97 | 2.29 | 1.75(1.74-1.77) | 1.080 |
| mRFP | NaCl | 0.1 (M) | 0.96 | 1.29 | 1.74 | 1.93 | 1.75(1.74-1.77) | 1.045 |
| mRFP | NaCl | 0.2 (M) | 2.21 | 1.87 | 0.83 | 1.27 | 1.75(1.74-1.76) | 1.072 |
| mRFP | NaCl | 0.4 (M) | 2.20 | 1.62 | 0.19 | 2.83 | 1.78(1.76-1.81) | 1.130 |
| mRFP | NaCl | 0.7 (M) | 3.02 | 1.83 | 0.76 | 1.15 | 1.73(1.73-1.74) | 1.055 |
| mRFP | NaCl | 1.0 (M) | 0.76 | 1.15 | 2.62 | 1.85 | 1.74(1.73-1.75) | 1.046 |
| mRFP | NaCl | 1.5 (M) | 2.14 | 1.54 | 0.83 | 2.18 | 1.76(1.75-1.79) | 1.108 |

| | | | | | | | | | |
|---|---|---|---|---|---|---|---|---|---|
| | mRFP | NaCl | 2.0 (M) | 2.27 | 1.47 | 1.19 | 2.13 | 1.75(1.74-1.78) | 1.064 |
| | mRFP | Trehalose | 0.0 (M) | 3.21 | 1.50 | 0.97 | 2.29 | 1.75(1.74-1.77) | 1.080 |
| | mRFP | Trehalose | 0.2 (M) | 0.55 | 0.74 | 5.74 | 1.74 | 1.70(1.69-1.70) | 1.131 |
| | mRFP | Trehalose | 0.4 (M) | 1.97 | 1.47 | 0.72 | 2.19 | 1.72(1.71-1.76) | 1.067 |
| | mRFP | Trehalose | 0.7 (M) | 1.35 | 1.30 | 1.31 | 1.97 | 1.70(1.67-1.72) | 1.018 |
| | mRFP | Trehalose | 1.0 (M) | 2.17 | 1.84 | 1.45 | 1.16 | 1.63(1.63-1.64) | 1.142 |
| | mRFP | Trehalose | 1.3 (M) | 0.83 | 0.74 | 4.99 | 1.67 | 1.60(1.60-1.61) | 1.131 |
| | mRFP | Glycerol | 0 (%) | 3.21 | 1.50 | 0.97 | 2.29 | 1.75(1.74-1.77) | 1.080 |
| | mRFP | Glycerol | 5 (%) | 1.76 | 1.58 | 0.24 | 2.70 | 1.79(1.77-1.87) | 1.056 |
| | mRFP | Glycerol | 10 (%) | 2.04 | 1.50 | 0.84 | 2.17 | 1.75(1.74-1.78) | 1.028 |
| | mRFP | Glycerol | 12 (%) | 2.11 | 1.59 | 0.15 | 3.04 | 1.76(1.74-1.79) | 1.078 |
| | mRFP | Glycerol | 14 (%) | 0.50 | 0.94 | 2.85 | 1.79 | 1.72(1.71-1.73) | 1.005 |
| | mRFP | Glycerol | 16 (%) | 1.96 | 1.59 | 0.11 | 3.30 | 1.77(1.73-1.80) | 1.058 |
| | mRFP | Glycerol | 20 (%) | 0.46 | 2.37 | 2.00 | 1.53 | 1.75(1.74-1.78) | 1.132 |
| | mRFP | Glycerol | 25 (%) | 0.59 | 2.38 | 2.38 | 1.52 | 1.76(1.74-1.80) | 1.149 |
| | mRFP | Glycerol | 30 (%) | 1.09 | 1.26 | 1.58 | 1.93 | 1.72(1.71-1.74) | 1.073 |
| | mRFP | Glycerol | 40 (%) | 0.80 | 0.80 | 4.58 | 1.74 | 1.67(1.67-1.67) | 1.044 |
| | mRFP | Glycerol | 50 (%) | 0.67 | 0.95 | 2.57 | 1.78 | 1.68(1.67-1.68) | 1.019 |
| | mRFP | PEG 3,350 | 0 (% FVO) | 3.21 | 1.50 | 0.97 | 2.29 | 1.75(1.74-1.77) | 1.080 |
| | mRFP | PEG 3,350 | 2 (% FVO) | 0.55 | 0.94 | 3.67 | 1.80 | 1.74(1.74-1.75) | 1.038 |
| | mRFP | PEG 3,350 | 7 (% FVO) | 3.14 | 1.85 | 0.80 | 1.10 | 1.75(1.74-1.76) | 1.015 |
| | mRFP | PEG 3,350 | 15 (% FVO) | 0.58 | 0.77 | 6.56 | 1.76 | 1.72(1.72-1.73) | 1.280 |
| | mRFP | PEG 3,350 | 20 (% FVO) | 0.75 | 0.91 | 3.89 | 1.78 | 1.70(1.70-1.71) | 1.048 |
| | mRFP | PEG 3,350 | 30 (% FVO) | 0.92 | 0.78 | 6.54 | 1.76 | 1.70(1.70-1.70) | 1.133 |
| | mRFP | PEG 3,350 | 40 (% FVO) | 1.34 | 0.91 | 4.66 | 1.80 | 1.69(1.69-1.69) | 1.160 |
| | mRFP | PEG 3,350 | 50 (% FVO) | 2.76 | 0.59 | 1.78 | 1.75 | 1.35(1.34-1.36) | 1.262 |
| | mRFP | PEG 6,000 | 0 (% FVO) | 3.21 | 1.50 | 0.97 | 2.29 | 1.75(1.74-1.77) | 1.080 |
| | mRFP | PEG 6,000 | 2 (% FVO) | 0.67 | 0.86 | 7.00 | 1.77 | 1.73(1.73-1.74) | 1.254 |
| | mRFP | PEG 6,000 | 7 (% FVO) | 1.18 | 1.23 | 2.71 | 1.88 | 1.74(1.73-1.76) | 1.028 |
| | mRFP | PEG 6,000 | 15 (% FVO) | 0.90 | 0.94 | 6.86 | 1.77 | 1.72(1.71-1.72) | 1.272 |
| | mRFP | PEG 6,000 | 20 (% FVO) | 1.07 | 0.88 | 4.90 | 1.80 | 1.71(1.71-1.72) | 1.004 |
| | mRFP | PEG 6,000 | 30 (% FVO) | 6.13 | 1.77 | 1.22 | 0.77 | 1.69(1.69-1.70) | 1.058 |
| | mRFP | PEG 6,000 | 40 (% FVO) | 1.57 | 1.87 | 1.43 | 0.86 | 1.57(1.57-1.58) | 1.145 |
| | mRFP | PEG 6,000 | 50 (% FVO) | 0.32 | 1.75 | 1.87 | 0.58 | 0.98(0.96-1.00) | 1.408 |
| | mRFP | PEG 10,000 | 0 (% FVO) | 3.21 | 1.50 | 0.97 | 2.29 | 1.75(1.74-1.77) | 1.080 |
| | mRFP | PEG 10,000 | 2 (% FVO) | 1.45 | 1.38 | 1.29 | 2.03 | 1.75(1.74-1.78) | 1.060 |
| | mRFP | PEG 10,000 | 7 (% FVO) | 2.97 | 1.87 | 1.02 | 1.17 | 1.74(1.74-1.76) | 1.039 |
| | mRFP | PEG 10,000 | 15 (% FVO) | 0.58 | 0.80 | 4.63 | 1.76 | 1.71(1.71-1.72) | 1.106 |
| | mRFP | PEG 10,000 | 20 (% FVO) | 0.82 | 0.86 | 4.79 | 1.78 | 1.70(1.70-1.71) | 1.078 |
| | mRFP | PEG 10,000 | 30 (% FVO) | 1.60 | 0.93 | 3.93 | 1.84 | 1.68(1.68-1.69) | 1.033 |
| | mRFP | PEG 10,000 | 40 (% FVO) | 2.84 | 0.81 | 2.28 | 1.81 | 1.45(1.45-1.46) | 1.124 |
| | mRFP | PEG 10,000 | 50 (% FVO) | 2.78 | 0.53 | 0.25 | 1.65 | 0.78(0.76-0.80) | 1.319 |
| | eGFP | NaCl | 0.0 (M) | 1.36 | 2.40 | 0.23 | 3.65 | 2.66 (2.65-2.67) | 1.296 |
| | eGFP | NaCl | 0.05 (M) | 0.21 | 1.67 | 1.02 | 2.75 | 2.64 (2.60-2.65) | 1.132 |
| | eGFP | NaCl | 0.1 (M) | 1.56 | 2.67 | 0.11 | 1.38 | 2.62 (2.62-2.63) | 1.469 |
| | eGFP | NaCl | 0.2 (M) | 0.07 | 4.13 | 0.65 | 2.45 | 2.68 (2.65-2.68) | 1.132 |
| | eGFP | NaCl | 0.4 (M) | 0.65 | 2.39 | 0.13 | 3.58 | 2.66 (2.65-2.67) | 1.136 |
| | eGFP | NaCl | 0.7 (M) | 0.55 | 2.47 | 0.02 | 5.33 | 2.70 (2.67-2.72) | 1.094 |
| | eGFP | NaCl | 1.0 (M) | 0.59 | 2.40 | 0.07 | 3.89 | 2.65 (2.64-2.66) | 1.113 |
| | eGFP | NaCl | 1.5 (M) | 0.69 | 2.36 | 0.14 | 3.44 | 2.61 (2.60-2.62) | 1.187 |
| | eGFP | NaCl | 2.0 (M) | 0.59 | 2.36 | 0.04 | 4.38 | 2.62 (2.60-2.62) | 1.033 |
| | eGFP | Trehalose | 0.0 (M) | 1.36 | 2.40 | 0.23 | 3.65 | 2.66 (2.65-2.67) | 1.296 |
| | eGFP | Trehalose | 0.2 (M) | 0.48 | 2.05 | 0.63 | 2.92 | 2.62 (2.61-2.64) | 1.005 |
| | eGFP | Trehalose | 0.4 (M) | 0.16 | 3.57 | 0.80 | 2.32 | 2.61 (2.60-2.65) | 1.075 |
| | eGFP | Trehalose | 0.7 (M) | 0.68 | 2.18 | 0.26 | 3.17 | 2.53 (2.51-2.55) | 1.042 |
| | eGFP | Trehalose | 1.0 (M) | 0.38 | 1.56 | 0.87 | 2.63 | 2.41 (2.41-2.42) | 1.070 |
| | eGFP | Trehalose | 1.3 (M) | 0.38 | 1.33 | 1.6 | 2.48 | 2.35 (2.35-2.35) | 1.029 |
| | eGFP | Glycerol | 0 (%) | 1.36 | 2.40 | 0.23 | 3.65 | 2.66 (2.65-2.67) | 1.296 |
| | eGFP | Glycerol | 5 (%) | 1.05 | 2.72 | 0.18 | 1.68 | 2.61 (2.61-2.62) | 1.142 |
| | eGFP | Glycerol | 10 (%) | 1.70 | 2.61 | 0.09 | 1.22 | 2.57 (2.57-2.57) | 1.333 |
| | eGFP | Glycerol | 12 (%) | 1.77 | 2.60 | 0.11 | 1.24 | 2.56 (2.56-2.57) | 1.394 |
| | eGFP | Glycerol | 14 (%) | 1.13 | 2.65 | 0.18 | 1.59 | 2.56 (2.55-2.56) | 1.084 |
| | eGFP | Glycerol | 16 (%) | 2.10 | 2.58 | 0.13 | 1.22 | 2.54 (2.54-2.55) | 1.445 |
| | eGFP | Glycerol | 20 (%) | 2.37 | 2.54 | 0.13 | 0.93 | 2.50 (2.50-2.51) | 1.574 |
| | eGFP | Glycerol | 25 (%) | 2.18 | 2.51 | 0.16 | 0.92 | 2.47 (2.47-2.48) | 1.344 |
| | eGFP | Glycerol | 30 (%) | 2.43 | 2.48 | 0.15 | 0.84 | 2.45 (2.45-2.45) | 1.505 |
| | eGFP | Glycerol | 40 (%) | 2.09 | 2.44 | 0.15 | 0.97 | 2.40 (2.40-2.41) | 1.239 |
| | eGFP | Glycerol | 50 (%) | 1.93 | 2.43 | 0.24 | 1.15 | 2.36 (2.36-2.37) | 1.095 |
| | eGFP | PEG 3,350 | 0 (% FVO) | 1.36 | 2.40 | 0.23 | 3.65 | 2.66 (2.65-2.67) | 1.296 |
| | eGFP | PEG 3,350 | 2 (% FVO) | 0.56 | 2.34 | 0.15 | 3.50 | 2.67 (2.65-2.70) | 1.063 |
| | eGFP | PEG 3,350 | 7 (% FVO) | 0.15 | 1.13 | 1.53 | 2.59 | 2.53 (2.53-2.54) | 1.096 |
| | eGFP | PEG 3,350 | 15 (% FVO) | 0.05 | 1.70 | 0.19 | 2.72 | 2.57 (2.55-2.61) | 1.020 |
| | eGFP | PEG 3,350 | 20 (% FVO) | 0.13 | 1.14 | 1.79 | 2.55 | 2.51 (2.50-2.51) | 1.222 |
| | eGFP | PEG 3,350 | 30 (% FVO) | 0.18 | 0.86 | 2.85 | 2.47 | 2.44 (2.43-2.44) | 1.523 |
| | eGFP | PEG 3,350 | 40 (% FVO) | 0.34 | 0.93 | 2.52 | 2.28 | 2.21 (2.20-2.21) | 1.525 |
| | eGFP | PEG 3,350 | 50 (% FVO) | 0.32 | 0.78 | 3.49 | 2.31 | 2.27 (2.26-2.27) | 1.606 |

| | | | | | | | | |
|---|---|---|---|---|---|---|---|---|
| eGFP | PEG 6,000 | 0 (% FVO) | 1.36 | 2.40 | 0.23 | 3.65 | 2.66 (2.65-2.67) | 1.296 |
| eGFP | PEG 6,000 | 2 (% FVO) | 0.69 | 2.31 | 0.26 | 3.30 | 2.66 (2.65-2.67) | 1.059 |
| eGFP | PEG 6,000 | 7 (% FVO) | 0.30 | 1.52 | 2.22 | 2.69 | 2.61 (2.60-2.61) | 1.131 |
| eGFP | PEG 6,000 | 15 (% FVO) | 0.18 | 0.99 | 3.30 | 2.58 | 2.55 (2.55-2.55) | 1.437 |
| eGFP | PEG 6,000 | 20 (% FVO) | 0.26 | 0.75 | 4.40 | 2.54 | 2.50 (2.50-2.51) | 1.640 |
| eGFP | PEG 6,000 | 30 (% FVO) | 0.28 | 0.80 | 4.21 | 2.45 | 2.42 (2.41-2.42) | 1.635 |
| eGFP | PEG 6,000 | 40 (% FVO) | 0.31 | 0.88 | 3.30 | 2.40 | 2.35 (2.35-2.35) | 1.393 |
| eGFP | PEG 6,000 | 50 (% FVO) | 0.33 | 1.03 | 1.38 | 2.24 | 2.12 (2.11-2.12) | 1.252 |
| eGFP | PEG 10,000 | 0 (% FVO) | 1.36 | 2.40 | 0.23 | 3.65 | 2.66 (2.65-2.67) | 1.296 |
| eGFP | PEG 10,000 | 2 (% FVO) | 0.13 | 1.20 | 1.99 | 2.65 | 2.61 (2.61-2.61) | 1.302 |
| eGFP | PEG 10,000 | 7 (% FVO) | 2.73 | 2.61 | 0.13 | 0.91 | 2.58 (2.58-2.62) | 1.381 |
| eGFP | PEG 10,000 | 15 (% FVO) | 3.93 | 2.57 | 0.22 | 0.76 | 2.54 (2.54-2.55) | 1.656 |
| eGFP | PEG 10,000 | 20 (% FVO) | 3.94 | 2.52 | 0.25 | 0.65 | 2.49 (2.49-2.50) | 1.729 |
| eGFP | PEG 10,000 | 30 (% FVO) | 3.10 | 2.48 | 0.23 | 0.56 | 2.45 (2.44-2.45) | 1.659 |
| eGFP | PEG 10,000 | 40 (% FVO) | 3.26 | 2.42 | 0.26 | 0.85 | 2.38 (2.37-2.39) | 1.517 |
| eGFP | PEG 10,000 | 50 (% FVO) | 0.31 | 0.99 | 1.98 | 2.27 | 2.19 (2.19-2.19) | 1.282 |
| mCitrine | NaCl | 0.0 (M) | 0.25 | 3.32 | 0.74 | 3.33 | 3.32(3.32-3.35) | 1.101 |
| mCitrine | NaCl | 0.1 (M) | 0.01 | 0.01 | 1.07 | 3.35 | 3.34(3.35-3.45) | 1.263 |
| mCitrine | NaCl | 0.2 (M) | 0.21 | 2.65 | 0.94 | 3.44 | 3.32(3.16-3.34) | 1.060 |
| mCitrine | NaCl | 0.4 (M) | 0.02 | 0.01 | 1.12 | 3.32 | 3.32(3.31-3.32) | 1.167 |
| mCitrine | NaCl | 0.7 (M) | 0.10 | 2.20 | 1.12 | 3.35 | 3.29(3.20-3.30) | 1.084 |
| mCitrine | NaCl | 1.0 (M) | 0.01 | 0.01 | 1.31 | 3.25 | 3.25(3.23-3.25) | 1.108 |
| mCitrine | NaCl | 1.5 (M) | 0.15 | 1.81 | 1.02 | 3.32 | 3.21(2.97-3.26) | 1.078 |
| mCitrine | NaCl | 2.0 (M) | 0.19 | 2.02 | 0.87 | 3.27 | 3.12(2.99-3.18) | 1.178 |
| mCitrine | Trehalose | 0.0 (M) | 0.25 | 3.32 | 0.74 | 3.33 | 3.32(3.32-3.35) | 1.101 |
| mCitrine | Trehalose | 0.2 (M) | 0.01 | 0.01 | 1.04 | 3.28 | 3.28(3.25-3.32) | 1.048 |
| mCitrine | Trehalose | 0.4 (M) | 0.11 | 3.24 | 1.00 | 3.24 | 3.24(3.24-3.37) | 1.114 |
| mCitrine | Trehalose | 0.7 (M) | 0.04 | 2.40 | 1.11 | 3.23 | 3.21(3.21-3.31) | 1.105 |
| mCitrine | Trehalose | 1.0 (M) | 0.36 | 2.80 | 1.20 | 3.16 | 3.08(3.08-3.17) | 1.108 |
| mCitrine | Trehalose | 1.3 (M) | 0.94 | 3.06 | 0.01 | 3.03 | 3.06(3.06-3.24) | 1.107 |
| mCitrine | Glycerol | 0 (%) | 0.25 | 3.32 | 0.74 | 3.33 | 3.32(3.32-3.35) | 1.101 |
| mCitrine | Glycerol | 5 (%) | 0.01 | 9.43 | 1.08 | 3.07 | 3.32(3.25-3.38) | 1.060 |
| mCitrine | Glycerol | 10 (%) | 0.01 | 9.68 | 1.11 | 3.05 | 3.28(3.21-3.51) | 1.050 |
| mCitrine | Glycerol | 12 (%) | - | - | - | - | - | - |
| mCitrine | Glycerol | 14 (%) | - | - | - | - | - | - |
| mCitrine | Glycerol | 16 (%) | 0.02 | 7.34 | 1.19 | 2.99 | 3.14(3.11-3.15) | 1.099 |
| mCitrine | Glycerol | 20 (%) | 0.01 | 9.63 | 1.24 | 2.95 | 3.17(3.11-3.19) | 1.090 |
| mCitrine | Glycerol | 25 (%) | 0.01 | 11.86 | 1.13 | 2.94 | 3.17(3.11-3.17) | 1.016 |
| mCitrine | Glycerol | 30 (%) | - | - | - | - | - | - |
| mCitrine | Glycerol | 40 (%) | 0.07 | 4.56 | 1.80 | 2.80 | 2.90(2.87-2.99) | 1.144 |
| mCitrine | Glycerol | 50 (%) | 0.01 | 9.47 | 1.55 | 2.83 | 2.90(2.85-2.91) | 1.185 |
| mCitrine | PEG 3,350 | 0 (% FVO) | 0.25 | 3.32 | 0.74 | 3.33 | 3.32(3.32-3.35) | 1.101 |
| mCitrine | PEG 3,350 | 2 (% FVO) | 0.36 | 3.39 | - | - | 3.39(3.35-3.40) | 1.041 |
| mCitrine | PEG 3,350 | 7 (% FVO) | 0.10 | 2.22 | 0.77 | 3.37 | 3.28(3.27-3.29) | 1.062 |
| mCitrine | PEG 3,350 | 15 (% FVO) | - | - | - | - | - | - |
| mCitrine | PEG 3,350 | 20 (% FVO) | 0.03 | 1.34 | 0.91 | 3.24 | 3.21(3.20-3.22) | 1.009 |
| mCitrine | PEG 3,350 | 30 (% FVO) | 0.05 | 1.63 | 0.98 | 3.14 | 3.10(3.09-3.10) | 1.063 |
| mCitrine | PEG 3,350 | 40 (% FVO) | 0.29 | 3.36 | 0.77 | 2.25 | 2.65(2.60-2.68) | 1.125 |
| mCitrine | PEG 3,350 | 50 (% FVO) | 0.91 | 2.20 | 0.19 | 3.27 | 2.45(2.41-2.53) | 1.120 |
| mCitrine | PEG 6,000 | 0 (% FVO) | 0.25 | 3.32 | 0.74 | 3.33 | 3.32(3.32-3.35) | 1.101 |
| mCitrine | PEG 6,000 | 2 (% FVO) | 0.00 | 0.01 | 0.88 | 3.33 | 3.33(3.22-3.33) | 1.172 |
| mCitrine | PEG 6,000 | 7 (% FVO) | 0.01 | 3.32 | 0.80 | 3.32 | 3.32(3.32-3.32) | 1.171 |
| mCitrine | PEG 6,000 | 15 (% FVO) | 0.19 | 3.23 | 0.86 | 3.23 | 3.23(3.23-3.45) | 1.128 |
| mCitrine | PEG 6,000 | 20 (% FVO) | - | - | - | - | - | - |
| mCitrine | PEG 6,000 | 30 (% FVO) | 0.76 | 2.59 | 0.17 | 3.74 | 2.87(2.86-2.90) | 1.072 |
| mCitrine | PEG 6,000 | 40 (% FVO) | 0.61 | 2.54 | 0.01 | 7.54 | 2.79(2.74-2.89) | 1.072 |
| mCitrine | PEG 6,000 | 50 (% FVO) | 0.01 | 0.01 | 1.47 | 2.46 | 2.46(2.45-2.52) | 1.149 |
| mCitrine | PEG 10,000 | 0 (% FVO) | 0.25 | 3.32 | 0.74 | 3.33 | 3.32(3.32-3.35) | 1.101 |
| mCitrine | PEG 10,000 | 2 (% FVO) | 0.05 | 2.01 | 0.81 | 3.40 | 3.35(3.34-3.36) | 1.019 |
| mCitrine | PEG 10,000 | 7 (% FVO) | 0.68 | 3.17 | 0.04 | 4.93 | 3.32(3.30-3.35) | 1.055 |
| mCitrine | PEG 10,000 | 15 (% FVO) | 0.36 | 3.22 | 1.18 | 3.22 | 3.22(3.20-3.22) | 1.197 |
| mCitrine | PEG 10,000 | 20 (% FVO) | - | - | - | - | - | - |
| mCitrine | PEG 10,000 | 30 (% FVO) | 0.66 | 2.30 | 0.38 | 3.37 | 2.79(2.78-2.80) | 1.011 |
| mCitrine | PEG 10,000 | 40 (% FVO) | 0.92 | 2.72 | 0.48 | 1.97 | 2.51(2.36-2.54) | 1.145 |
| mCitrine | PEG 10,000 | 50 (% FVO) | 1.17 | 2.62 | 0.75 | 1.15 | 2.29(2.15-2.70) | 1.111 |

**Supplementary Table 2**: a list of the residues exhibiting a network of contacts with the fluorophore of mCherry in the MD simulations

| Residue | Probability (%) at 0% FVO PEG | Probability (%) at 42% FVO PEG |
|---|---|---|
| F14 | 84.7 | 77.6 |
| Q42 | 99.9 | 99.5 |
| T43 | 88.3 | 75.7 |
| A44 | 99.8 | 99.5 |
| L46 | 6.3 | 8.7 |
| S62 | 16.8 | 24.6 |
| P63 | 99.9 | 100.0 |
| Q64 | 87.2 | 84.9 |
| F65 | 100.0 | 100.0 |
| S69 | 100.0 | 100.0 |
| K70 | 65.7 | 63.4 |
| F91 | 3.1 | 4.3 |
| W93 | 99.3 | 98.6 |
| R95 | 100.0 | 100.0 |
| Q109 | 49.9 | 51.6 |
| W143 | 31.7 | 29.0 |
| E144 | 2.4 | 1.5 |
| A145 | 15.3 | 17.2 |
| S146 | 41.1 | 47.8 |
| E148 | 4.2 | 7.1 |
| I161 | 95.4 | 99.4 |
| Q163 | 85.7 | 83.0 |
| V177 | 0.7 | 1.4 |
| Y181 | 29.0 | 37.4 |
| I197 | 99.5 | 99.6 |
| K198 | 3.7 | 1.7 |
| L199 | 100.0 | 100.0 |
| Q213 | 99.7 | 99.8 |
| Y214 | 57.1 | 49.2 |
| E215 | 100.0 | 100.0 |
